# Heat stress impacts the gut microbiome of Atlantic salmon by promoting growth of Vibrionaceae and is associated with extensive cast production

**DOI:** 10.1101/2024.11.23.625002

**Authors:** John P. Bowman

## Abstract

Atlantic salmon (*Salmo salar*) when exposed to heat stress have reduced voluntary feeding and exhibit changes in digesta consistency. This study was performed to determine what bacterial species occur and the overall bacterial abundance in the gut microbiome in heat stressed Atlantic salmon. For this Atlantic salmon in seawater tanks at 15°C were fed for 2 to 4 weeks. Tank temperatures were increased to 19°C until voluntary feeding abated at which point tank temperatures were cooled to 15°C for 4 weeks. At the end of each temperature phase the fish were stripped of feces and microbiome profiles were determined using 16S rRNA V1-V3 metabarcoding. The tank experiment was repeated three times in successive years. Vibrionaceae comprised most reads after the warm phase completed. The prominent levels of Vibrionaceae were accompanied by a large predominance of cast (sloughed intestinal mucosa) containing fecal samples. qPCR estimated Vibrionaceae cell populations increased 1.9-3.4 log units/g after the warm phase but and slightly decreased by 0.3-1.1 log units/g after the 15°C recovery phase. The results indicated heat stress induced inappetence corresponds to increased cast production accompanied by predominance of Vibrionaceae. Vibrionaceae colonizing the Atlantic salmon gut should be the focus of studies on the microbiology of thermally induced inappetence and dysbiosis in Atlantic salmon.

**Key Contribution:** The experiments confirm that thermal stress increased cast production in Atlantic salmon. The increase in cast production coincides with increased predominance of Vibrionaceae. Though the growth of Vibrionaceae in the gut of salmon is not associated with acute disease it may represent a dysbiosis effect. The results suggest extended summers in regions where sea surface temperatures are at the tolerance limits require increased husbandry management of farmed fish. In addition, more studies are needed to determine if novel approaches can limit overgrowth of Vibrionaceae.

## 1. Introduction

Farmed Atlantic salmon prefer water temperatures between 13 to 16°C [1]. The temperature limits and associated physiological impacts depend on the life stage [2] and genetics to a lesser degree [3]. Heat stress can significantly impact farmed salmon health, productivity and their subsequent marketability [4, 5]. The main sign of heat stress in Atlantic salmon and other salmonid species is lethargic behavior and reduced or no voluntary feeding [5]. This eventually results in substantial size and condition variation within cohorts. Lack of feeding may also lead to inconsistent flesh color due to uneven deposition of dietary astaxanthin into white muscle [6]. Heat stress and inappetence could be partly due to impacts on renal, liver and osmoregulatory functions. Summer conditions has also been associated with a steep increase in the excretion of pale yellow sloughed intestinal mucosal layer referred to as “pseudofeces” or fecal casts [7]. To date these issues are managed alongside other summer-related risks including amoebic gill disease [8], toxic algal blooms [9], sea lice infestation [10], and reduction in oxygen availability [11].

Fish have a gut microbiome that reflects their environment [12]. In farmed fish there is a requirement to decipher microbiome data to determine what bacterial taxa make up the active colonizing community. This avoids confounding data where bacteria are misinterpreted as being part of the microbiome. For example, members of the genus *Geobacillus*, an endospore-forming thermophile growing best at 60-70°C and unable to grow <35°C, can be readily detected in Atlantic salmon digesta samples [13]. This occurrence is consistent with bacterial cells or remnant DNA that arrive in the gut within feed-pellets [14]. In addition, substantial numbers of bacteria (and other seawater microorganisms) present in water imbibed by the fish passively transit through the gut [13]. Furthermore, salmon smolt, when transferred to sea pens, are physiologically adapted to the marine environment, however their gut microbiota still dynamically changes over time [15-17]. Diet may also influence the community composition of farmed Atlantic salmon though predominant bacterial species emerge given time [18]. A major autochthonous (colonizer) species present in wild and farmed Atlantic salmon is “Candidatus Mycoplasma salmoninarum” corrig. [19]. This unculturable species has been detected in Atlantic salmon gut microbiomes world-wide but is particularly predominant in salmon growing in high latitude [18]. Under more temperate conditions (sea surface temperatures of 15-20°C) surveys have shown that the gut bacteria include a predominance of Vibrionaceae (*Aliivibrio, Vibrio, Photobacterium*) especially after fish experience their first summer [7, 13]. High abundances of bacteria such as *Aliivibrio* have been suspected to be more predominant in fish expressing casts (also referred to as pseudofeces) and feces with a watery composition that may contain cast material [7, 17]. The connection of casts with the presence of Vibrionaceae to dysbiosis and other negative health impacts in Atlantic salmon [20] has yet to be shown conclusively. Nevertheless, Vibrionaceae are of interest to salmon health due to their potential to cause disease [21-23] and as potential beneficial bacteria [24].

Due to their predominance in farmed salmon during summer in warmer water salmon farming regions there is a need to understand whether Vibrionaceae have a role in thermally associated dysbiosis [25]. This includes understanding capabilities related to persistent colonization and in vivo functionality. At this stage there is no data on what specific gut microbiome bacterial species are promoted under elevated temperatures. It is assumed these would be Vibrionaceae but whether this includes one or few species or a broader diversity and whether repeated experiments provide reproducible responses has yet to be shown. In this study the first aim was to determine the effect of temperature on the Atlantic salmon gut microbiome. This involved assessment of the species composition in digesta samples and an assessment of fecal consistency of samples from fish reared in seawater tanks initially set at 15°C. For the experiments. Tank water temperature was raised to a range (18.5-19°C) that reduced voluntary feeding. The temperature used for heat stress was lower than extremes that have occurred near where the experiments took place [26] but potentially could eventually happen in the near future at temperate latitudes. The experiment was repeated three times in separate years to determine to understand the reproducibility of microbiome responses. The final goal was to measure to bacterial populations levels exist in Atlantic salmon reared under the temperature conditions of the experiment thus establishing the degree of bacterial proliferation.

## 2. Materials and Methods

### 2.1. Research Design

Tank temperature experiments were performed at the Experimental Aquaculture Facility (EAF) of the University of Tasmania, Taroona, Hobart. The EAF houses a multi-tank recirculating aquaculture system. In this experiment Atlantic salmon were exposed to staged temperature exposures as described in Figure 1. Temperature-based experiments were repeated three times, in successive years (2016, 2017, 2018 - from March-April to September). For sampling at the end of each of the temperature phases fish were obtained from multiple tanks.

**Figure 1.**
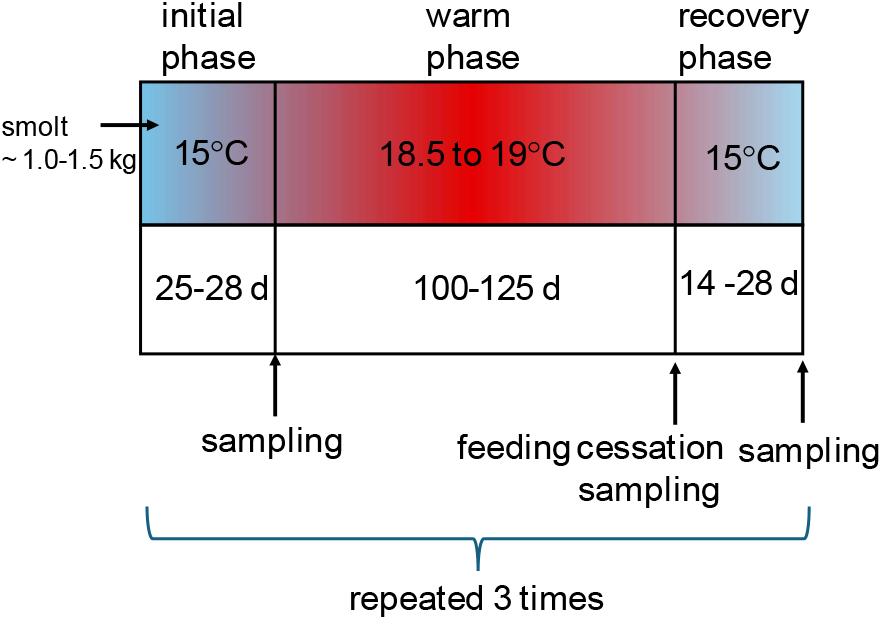
Research design for the heat stress experiment. Experiments were performed in seawater-containing tanks. Sampling involved stripping feces from random fish.

### 2.2. Tank set up and sampling

The tanks used were filled with 1 μm filtered seawater from the Derwent Estuary at a daily exchange rate of 10% volume. Water quality was maintained using a combination of protein skimmers and a series of treatment steps including <40 μm drum filtration, ammonia removal via nitrification-based biofilters, and UV light irradiation. For the experiments, the operating conditions used were like that described by Foddai et al. [27]. Smolt, already adapted to seawater, were transferred to a 13000 L seawater tank and fed until they reached an average weight of approximately 1.5 kg. Fish were then evenly distributed between twelve 7000 L tanks in the common RAS system with a stocking density of 50-100 fish per tank and photoperiod set at 12:12 light: dark hours (illumination at 150□±□50□μmol). The tanks were maintained at 15°C for 25 to 28 d at which point fish were randomly chosen for sampling. Tank temperatures were then increased at 0.1°C per day over 7 weeks to reach on average 18.5-19.0°C and maintained at this temperature for at least 100 to 125 d at which point fish ceased feeding. Tank temperatures were then reduced to 15-16°C and the fish sampled after 2 to 4 weeks. Dissolved oxygen was maintained at 100□±□2% saturation, water salinity at 35 psu and pH at 7.8 ±□0.1. Fish were hand-fed a post smolt commercial diet continually throughout to maintain satiation.

### 2.3. Sampling procedures

Fish were sampled randomly from tanks by using a dip-net. For the initial time point 10, 10, and 24 fish were sampled in the three trials, respectively. After the warm water phase typically 6 fish were sampled from each tank resulting in 60, 42 and 60 fish being sampled in the three trials, respectively. In the recovery 15°C phase 25, 36, and 12 fish were sampled in each of the three trials, respectively.

Captured fish were transferred into aerated tubs containing 17 ppm Aqui-S. For fecal stripping, ventral surfaces of anaesthetized fish were wiped with ethanol to minimize contamination, and samples were obtained by gently pinching and massaging the fish from the midline above the pelvic fins down towards the vent; 0.5 to 1 g of digesta was aseptically collected and placed into a sterile 15 ml tube. The tubes were snap frozen in a dewar containing liquid nitrogen or placed on ice. Fish were returned to tanks upon recovery from anesthesia. Frozen tubes were transported back to the laboratory and stored at – 80 °C until DNA was extracted. All fish handling procedures are covered under University of Tasmania Animal Ethics Permit A0015452.

### 2.4 Fecal scoring

The consistency of the fecal samples and presence of fecal casts was achieved using a categorical scheme of scoring method [17]. For this the fecal scores range from 1 to 5 and are described as follows: 1 - firm digesta, no casts; 2 – digesta with mid-range water content, no casts; 3 – digesta watery, no casts; 3.5 - digesta watery and containing pale yellow or white cast material; 4 – only cast material obtained; 5 – empty gut, with only fluid with some mucus obtained.

### 2.5. DNA extraction

Total genomic DNA from 0.25 g stripped fecal samples were extracted using the QIAamp DNA Stool Kit (Qiagen, Crawley, United Kingdom) following the manufacturer’s specifications. Two or three subsamples from each fecal sample collected were extracted with the final DNA pooled. The concentration of DNA in extracts was estimated using fluorimetry using the Quant-iT PicoGreen dsDNA assay kit (Thermo Fisher Scientific). Samples that had a score of 5 for fecal consistency (13 of 274 total samples) were not extracted.

### 2.6. MiSeq Illumina-based 16S rRNA gene sequencing

All DNA amplifications and sequencing were conducted by using the Illumina MiSeq platform and 300 bp paired end sequencing chemistry. Amplicons were prepared using bacterial 16S V1-V3 rRNA gene primers V1-27F (5’-AGA GTT TGA TCM TGG CTC AG-3’) [28] and V3-519R (GWA TTA CCG CGG CKG CTG-3’) [29]. In each run no-template blank controls were used to account for kit and laboratory contamination. The first included buffer ATE (low-EDTA elution buffer) from the QIAamp Fast DNA Stool Mini Kit (Qiagen, Crawley, United Kingdom) from the University of Tasmania laboratory where DNA extraction occurred to account for laboratory contamination. In addition, a blank distilled water control was also used to account for contamination in the sequencing laboratory (Ramaciotti Centre for Genomics, Sydney, NSW, Australia).

### 2.7. Sequence data processing

Paired-end sequences were denoised, dereplicated, chimera filtered and merged using QIIME 2 implemented ‘denoise-paired’ command with DADA2 [30] to produce amplicon sequence variants. Default settings were used, and all reads with a median PHRED score <20 were discarded. Denoised data from all samples were then merged to form a single feature table for subsequent processing. To simplify the data OTUs were then clustered at 99% sequence identity threshold using cluster-features-de-novo in QIIME 2 and using VSEARCH [31]. Using QIIME 2, a phylogenetic tree was generated using align-to-tree-mafft-fasttree from the q2-phylogeny plugin (MAFFT for multiple sequence alignment and FastTree to generate a phylogenetic tree from a masked alignment [32]. Classification of OTUs was performed against the SILVA 138.1 [33] and NCBI curated prokaryote 16S rRNA gene (current to May 2024) databases. The R package Decontam identified 14 contaminant OTUs from the no template controls and these were removed from the sequence dataset [34].

### 2.8. Diversity analysis

For diversity estimates chloroplast and mitochondrial-derived 16S rRNA gene sequences were excluded from the dataset as well as any reads unassigned to bacterial phyla. The OTU table was imported into Primer 7 (Primer-E, Auckland, New Zealand) to calculate taxonomic diversity based on the Shannon index (H’).

### 2.9. Multivariate analysis

The filtered OTU read data at each time point and were summed and converted to centered log ratios (CLRs) [35] separately for each of the three repeat experiments. CLR data was imported into Primer 7 and a resemblance matrix was determined using Euclidean distance analysis [36]. In Primer 7 non-metric multidimensional scaling and analysis of similarity was used preliminarily examine the similarity and overlapping of sample types inclusive of time point. Canonical analysis of principal coordinate analysis (CAP,[37]) was also used to ascertain how distinct treatment-based groupings were by utilizing the cross-validation feature. The number of axes used to fit the data was constrained to the minimum number (m=4 to 10) based on the recommendations of Ratkowsky [38]. PERMANOVA [39] was used to determine the influence of treatment and fecal score. For this analysis, all raw data was used, assumed partial sum of squares, 9999 permutations and Monte Carlo adjustment of p-values.

### 2.10. Quantitative PCR

Quantitative PCR (qPCR) was used to estimate 16S rRNA gene copy numbers in the same DNA extracts used for microbiome profiling. The primers used were bacterial universal primers 1369F (5’-CGG TGA ATA CGT TCY CGG-3’) and 1492R (5’-GGW TAC CTT GTT ACG ACT T-3’) [40]. The SensiFAST SYBR, NO-ROX Kit (Bioline Australia) was used for all qPCR reactions. A 1369F/1492R amplicon was generated from *Aliivibrio finisterrensis* LMG 23869 of 151 bp size using the MyTaq Direct PCR kit (Bioline Australia) and utilizing a PTC-200 Thermal Cycler (MJ Research). The amplicon was checked using agarose gel electrophoresis and purified using 20 μL MagBio HighPrep PCR magnetic beads (MagBio Genomics, United States) with DNA eluted into pre-warmed (50°C) of 20 μL 10 mM Tris-HCl pH 8.0 supplemented with 50 mM NaCl. The Quant-iT PicoGreen dsDNA assay kit and was then used to adjust the amplicon concentrations to 50.5 ± 1.0 and 5.0 ± 0.6 ng/μL with 10 mM Tris-HCl pH 8.0. The amplicon solutions were then diluted in ten-fold steps with 10 mM Tris-HCl pH 8.0. The diluted amplicon solutions were utilized to relate 16S rRNA copy number to critical threshold (Ct) level. For the standard curve, gene copy numbers per unit amount of DNA was calculated using the equation:

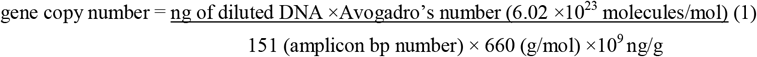

For the analysis 8 fecal DNA samples were analysed from each temperature treatment for each trial (total samples n=72). The composition of the qPCR 10 μL reaction mixtures were 5 μL of 2 × SYBR green reaction mix, 1 μL water, 1 μL (5 μM) forward primer, 1 μL (5 μM) reverse primer, and 2 μl DNA template. Reactions were performed in a Rotor-Gene Q instrument (QIAGEN, Hilden, Germany). Cycle profiles consisted of an initial hold at 95°C for 3 min, followed by 35 cycles of denaturation at 95 °C for 10 s annealing of primer at 60°C for 15 s, and extension at 72 °C for 20 s. Melt curve analysis, determination of PCR efficiency and coefficient of variation were determined [41]. For each tested Atlantic salmon fecal DNA sample Ct values were generated three -times within each qPCR run. Three separate qPCR runs were performed for each sample set.

### 2.11. Statistical analysis

Comparisons between diversity indices utilized the Kruskal-Wallis test for non-normal data. Welch’s t-test was used for pair-wise significance analysis with an α level of ≤0.01 considered significant. The designation “p(MC)” indicates the significance was adjusted by Monte-Carlo simulation.

## 3. Results

### 3.1. Fecal consistency

For salmon held at 15°C for 24 to 28 d in the three trials most samples had fecal consistency scores of 1 to 3 (75%, 90%, and 96% for the 3 trials, respectively) (Figure 2). Most digesta samples (68%, 83%, and 87%) from fish sampled at the end of the 18.5-19°C temperature (“warm”) phase contained fecal casts, represented by either score 3.5 or 4. The number of samples of each of these categories were approximately equal in number in trials 2 and 3, however only one sample consisted of entirely cast material from the trial 1 warm water phase (Figure 2). In samples collected after the 2 to 4-week 15°C “recovery” phase the proportion of fecal cast-containing samples was observed to vary between trials. These proportions were 30%, 78%, and 36% for the three trials, respectively (Figure 2).

**Figure 2.**
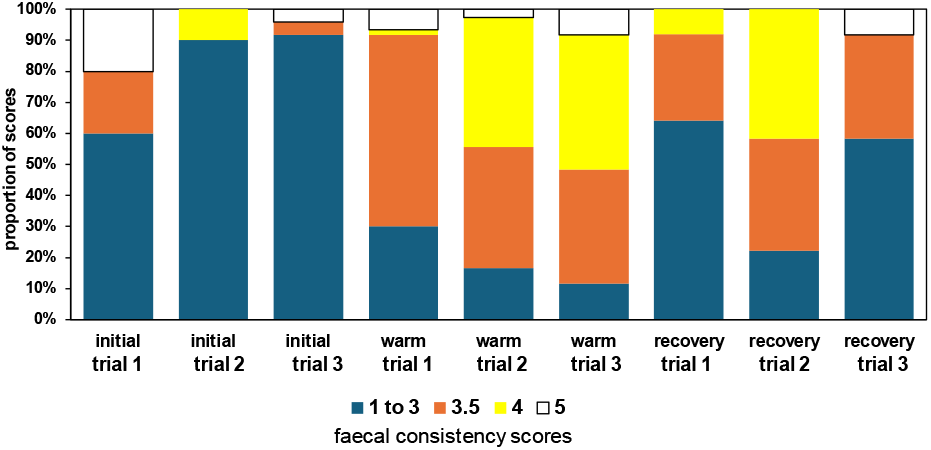
Fecal consistency scores of samples from Atlantic salmon that were exposed to different temperature treatments as defined in Figure 1 and in the Methods and Materials. Digesta only samples have scores of 1 to 3. Cast-containing samples are indicated by scores 3.5 and 4. Empty gut samples have a score of 5.

### 3.2. Microbiome diversity and structure between treatments and trials

The initial samples for all trials showed the highest H’ values (average H’ = 2.1 to 3.2). The warm temperature phase had much lower diversity (H’ = 0.2 to 0.8, p<0.0001) (Figure 3). The return to 15°C resulted in an increase in H’ values (H’ = 0.5 to 1.7, p<0.01) (Figure 3). In trial 1 the differences between the temperature treatments were the least obvious though there was a significant difference between the initial and warm phase H’ values (Figure 3, p<0.01). There was a significant effect on diversity with fecal consistency scores with scores 1 to 3 much higher in diversity (H’ = 2.2 to 3.0, p<0.0001) than that observed for either scores 3.5 (H’ = 0.3 to 0.6) or 4 (H’ = 0.1 to 0.3) (Figure 3).

**Figure 3.**
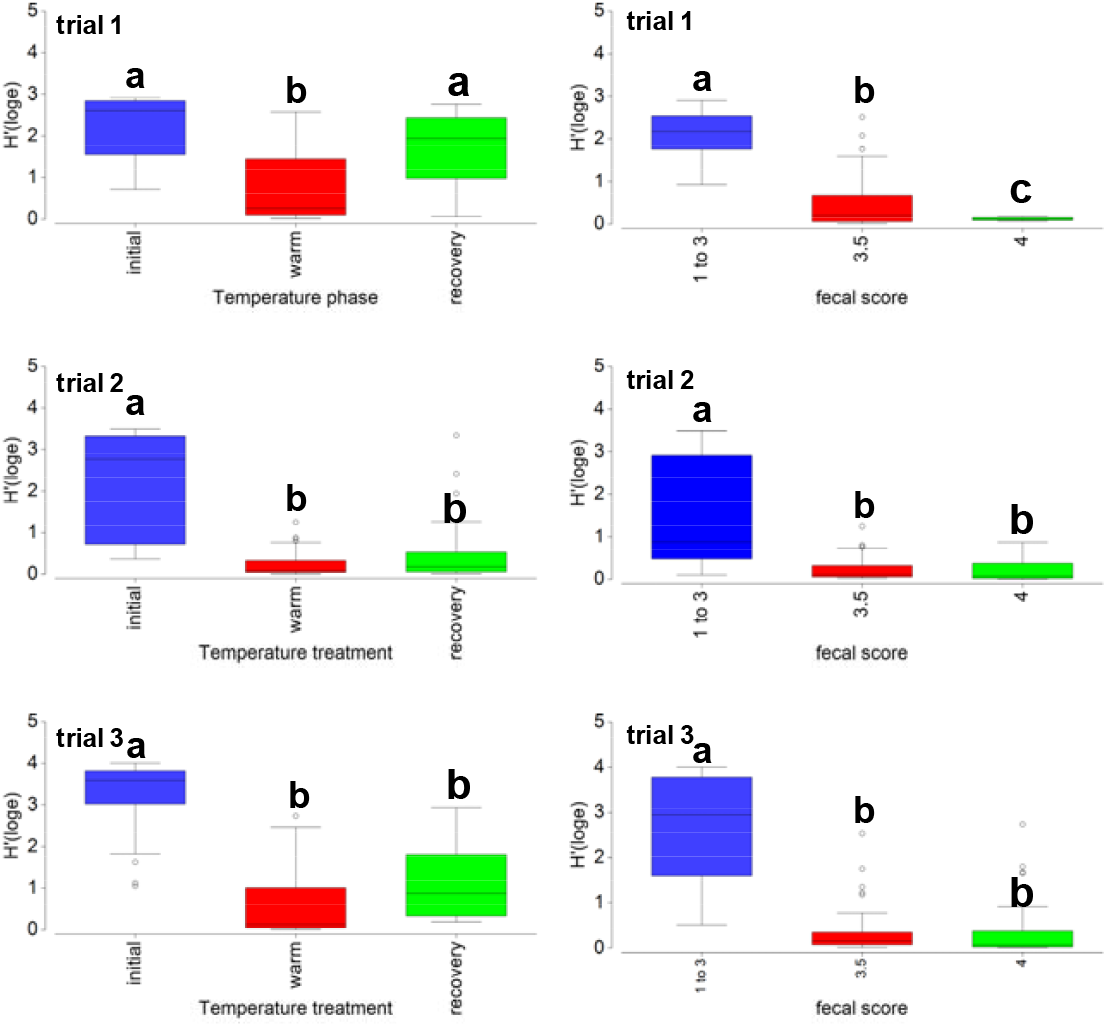
Shannon diversity (H’) of bacteria present in the fecal samples from Atlantic salmon exposed to different temperature treatments as defined in Figure 1 and in Methods and Materials. Fecal score designations are described in Figure 2 and in the Methods and Materials section. Letters above box plots that are different indicate H’ is significantly different (p<0.01) from data in the same graph.

PERMANOVA indicated in trials 1 and 3 the temperature treatment was significant on its effect on the bacterial community (pseudo-F values of 3.66 and 2.88, p(MC)<0.0001). CAP cross-validation (Figure 4) indicated the temperature treatment groups were clearly distinguished (80-100% correct classification). For trial 2 the p(MC)-value was 0.081 (pseudo-F =1.62). CAP cross-validation for trial 2 temperature treatment profiles was lower (67-80%) than the other trials. Fecal consistency scores had a significant influence on community structure overall in all three trials (pseudo-F = 2.53 to 6.80, p(MC)<0.0007) with samples with scores 1 to 3 well differentiated from the cast containing samples (CAP 85 to 97%, Figure 4).

**Figure 4.**
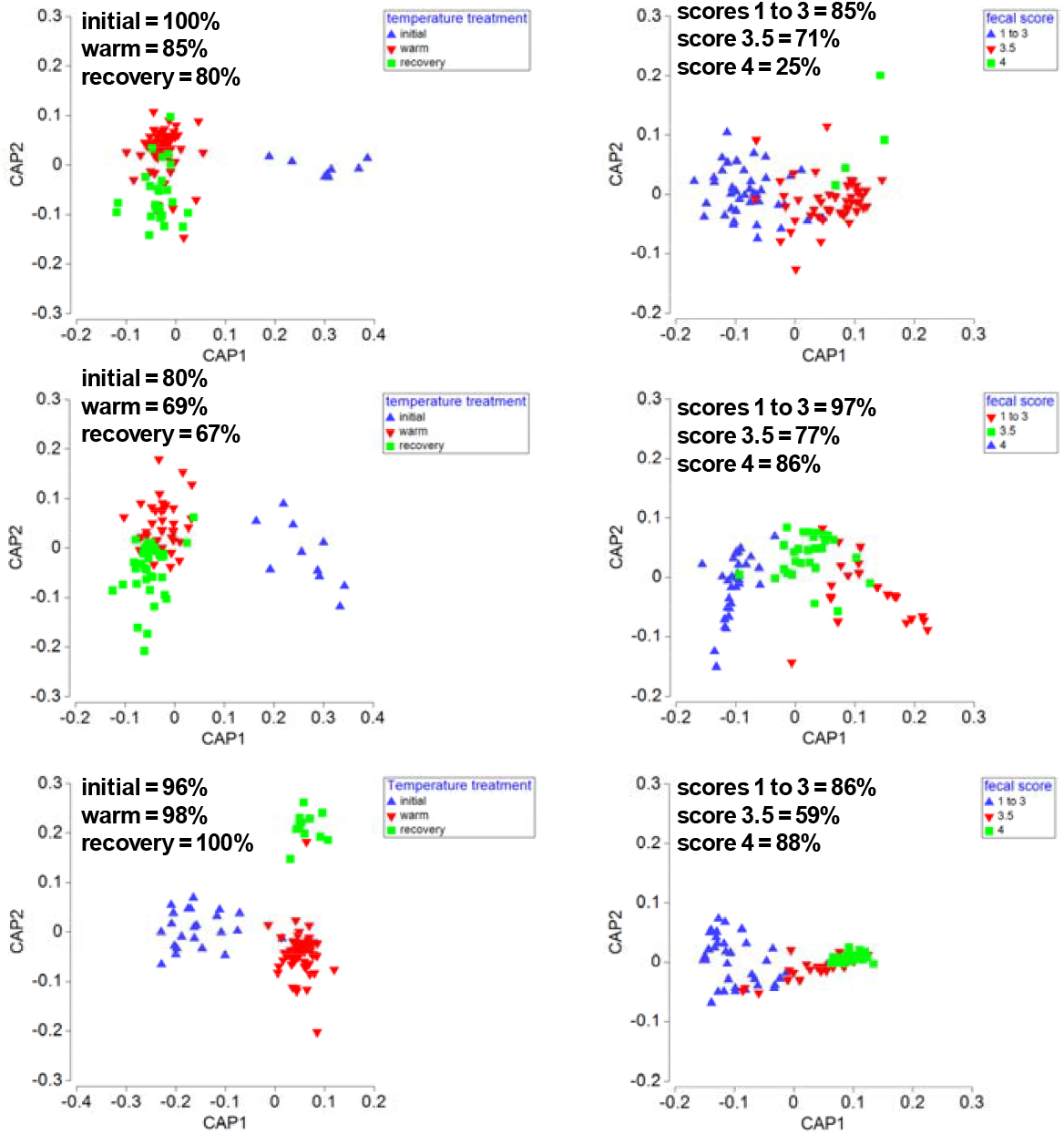
CAP plots comparing factors - temperature treatment (see Figure 1) and fecal scores (Figure 2). Data cross-validation results are summarised for each graph, indicating the proportion of samples that were successfully grouped according to the indicated factor. The lower the % classification the less defined the group is determined on the basis of correlation.

### 3.3. Bacterial composition

In the initial samples in the three trials most specimens analysed possess members of family Vibrionaceae including *Vibrio, Aliivibrio* and *Photobacterium* (totaling 24, 39 and 15% of total reads by trial order) and a high feed DNA background. The latter was indicated by the high proportions of plant-derived feed (wheat, lupin, faba bean, pea) chloroplast- and mitochondria-derived 16S rRNA reads (18, 31 and 47% of reads by trial order, Figure 5). *Vibrio* OTUs were predominant in trials 1 and 2 (24% and 39% of reads, Figure 5) at the initial stage while *Aliivibrio* or *Photobacterium* OTUs had initially low abundance (<0.03%, Figure 5). Unlike *Photobacterium, Aliivbrio* after the warm phase increased to about 1.6% of reads (CLR -2.1. to -0.3). In trial 3 the initial proportions of *Aliivibrio* and *Vibrio* were more equal (8.5 and 5.7%, respectively).

**Figure 5.**
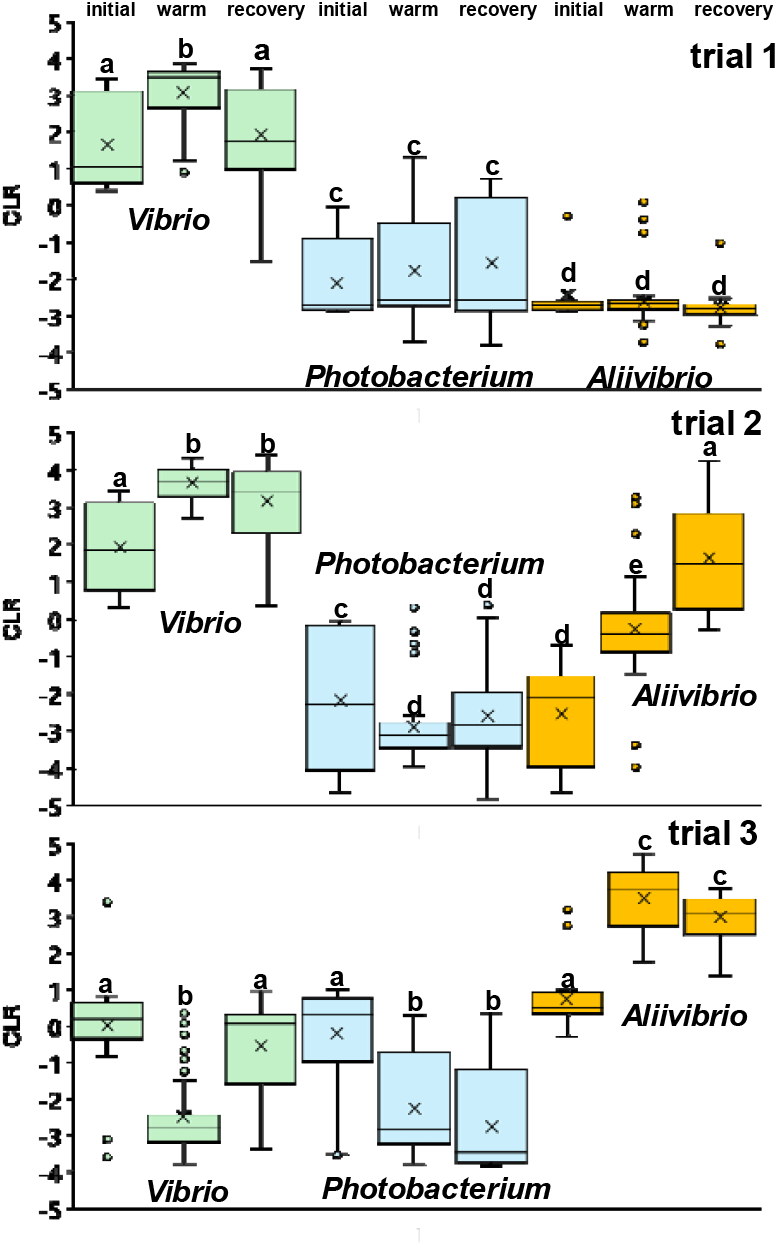
The caption for Figure 5 was revised for clarity as follows. “Figure 5. Comparisons of the relative abundance (CLR, centered log ratio) of Vibrionaceae from fecal samples of Atlantic salmon exposed to different temperatures. The temperature phases includes initial - 15°C (4 weeks), warm - 18.5-19°C (up to 100 d), recovery - 15°C (2 to 4 weeks). The data comes from three separate trials performed in different years. Letters above box plots that are different indicate that average CLR values were significantly different (p<0.01) to other box plot treatment data in the same trial.

The feed background was smaller in the warm phase temperature treatments with proportions of plant-derived reads dropping to 8.9, 3.8 and 1.5% in order of trials. The microbial compositions after the warm water phase differed each time and were dominated by either *Vibrio, Aliivibrio* or both genera (Figures 5 and 6). *Photobacterium* was always at low levels during the trials. In trials 1 and 2 Vibrio increased significantly in predominance making up 78 and 84% of total reads, respectively (Figure 5). A comparison using CLR values confirmed the increase was significant (p<0.0001, CLR averages 1.6 to 3.1 and 1.9 to 3.7, Figure 6), In trial 2 *Aliivibrio* also exhibited a significant abundance increase from a low level (0.004% of total reads) initial to 1.4% of total reads (CLR -2.6 to -0.3, Figure 6). In trial 3 *Aliivibrio* was instead the only predominant taxon (95% of reads, CLR -1.6 increasing to 3.5, Figure 6) with the presence of *Vibrio* dropping to low levels (0.005% of reads).

**Figure 6.**
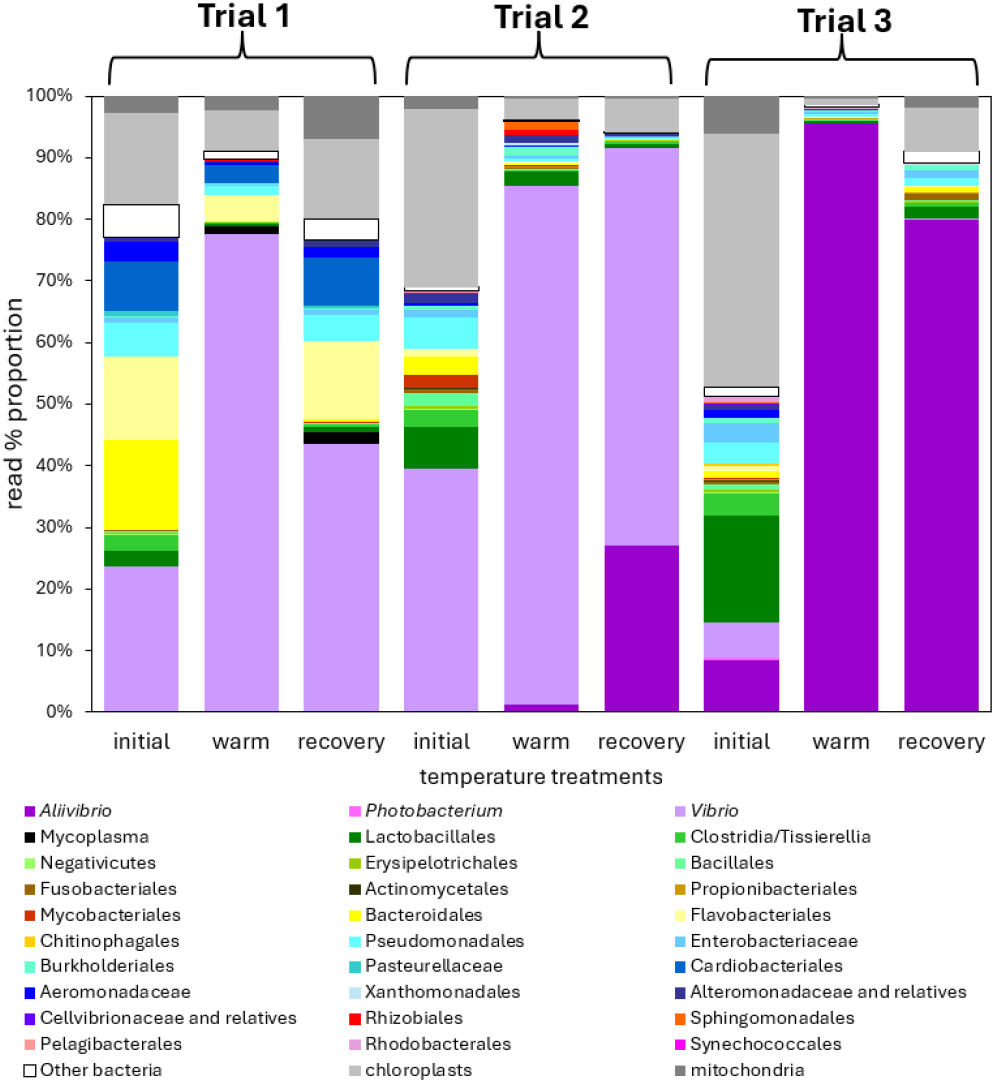
Proportional composition of bacterial communities from stripped fecal samples from Atlantic salmon reared in seawater containing tanks and exposed to different temperatures. The temperature phases include: initial - 15°C (4 weeks), warm 18.5-19°C (up to 100 d), recovery - 15°C (2 to 4 weeks). The data comes from three separate trials performed in different years. Chloroplast and mitochondrial 16S rRNA reads are included since these components relate to the feed and the proportion of reads changes between the temperature treatments, especially in trials 2 and 3.

In the cooler recovery phase, the feed background became again more obvious increasing to 20, 6, and 9% of total reads in the order of the trials. The data suggests that overall Vibrionaceae proportions correspondingly declined slightly, except in trial 2. CLR comparisons indicated in trials 1 and 2 *Vibrio* amassed 44 and 64% of total reads, decreasing relative to the warm phase sample levels (CLR averages dropping 3.1 to 1.9, 3.6 to 3.1, Figure 6). In trial 2 the *Aliivibrio* read proportion exhibited an additional 20-fold increase to 27% (CLR average increase -0.3 to 1.6, Figure 6). In trial 3 *Aliivibrio* made up 80% of reads and was slightly reduced in relative abundance compared to the warm phase levels (CLR 3.5 to 3.1, Figure 6).

Reads attributed to other bacteria were detected in the samples and were found to have highest proportions in initial samples (Fig. 6). When the data is integrated across all three trials the more abundant of these included Lactobacillales (2.5-17.2% of total reads), Clostridia/Tisseriellia (2.6 to 3.5%), Bacteroidales (0.9 to 14.5%), Flavobacteriales (1.0 to 13.4%), Pseudomonadales (3.4 to 5.6%), and Aeromonadaceae (0.4 to 3.2%). All these groups became less abundant (reduced between 2 to 20-fold) in the warm water phase (Fig. 6). In the recovery phase most of these groups correspondingly became more abundant by 2 to 4-fold (Fig. 6). At the genus level there was observed considerable variability between trials in terms of relative abundance. Since non-Vibrionaceae bacterial OTUs are not promoted during heat stress the genus specific details are not discussed here. Detailed results at the genus level are provided in Supplementary Table S1.

Two OTUs most similar to *Vibrio scophthlami* made up all Vibrionaceae reads in trial 1 (99.8% of Vibrionaceae reads; 69.6% of total reads), while reads related to *Aliivibrio finisterrensis* made up all reads in trial 3 (99.2% of Vibrionaceae reads; 84.1% of total reads). In trial 2 most Aliivibrio reads were most closely related to *Aliivibrio sifiae* and related species while Vibrio reads mostly belonged to *Vibrio scophthalmi* or a closely species. The taxa indicated were also the predominant taxa associated with samples with casts in the respective trials, making up most reads therein (81.0 to 99.8% across all cast-only samples).

### 3.4. Quantitative PCR

The standard deviation in estimated total copy number between the sets of 8 samples for each treatment and trial was 0.4 to 1.1 log units with PCR efficiency (E) averaging 0.92 ± 0.6 and coefficient of variance ranged between 17 to 51% between separate runs. The initial samples averaged log 16S rRNA gene copy number per g from 8.5 to 9.0 (Table 1). The copy number abundances increased 1.0, 1.0, and 1.9 log copies/g (to 9.5 to 10.4 log copies/g) in the warm phases of the respective trials for samples containing only digesta and for samples containing digesta and casts (Table 1). Cast-only samples had only slightly lower abundances (Table 1) as measured for samples from trials 2 and 3. The recovery phase samples had similar 16S rRNA gene copy concentrations to the warm phase ranging from 9.2 to 9.7 log copies/g. Overall, there was a significant increase in the abundance of 16S rRNA gene copies between the initial and warm phases (p<0.008) for all trials. The differences between warm and recovery phase were only marginally significant (p<0.03) for trial 3 samples. There also was no difference between samples containing digesta and casts and cast-only samples when compared for the warm or recovery phases in trials 2 and 3 (Table 1).

**Table 1.** Estimates of total 16S rRNA gene copies and Vibrionaceae in Atlantic salmon fecal samples.

|  | Total 16S rRNA<br>log copies/g |  |  | Estimated Vibrionaceae log cells/g |  |  | Change<br>(warm –<br>initial)<sup>3</sup> | Change<br>(warm –<br>recovery)<sup>3</sup> |
| --- | --- | --- | --- | --- | --- | --- | --- | --- |
|  | Initial | Warm | Recovery | Initial | Warm | Recovery | Log unit change |  |
| Trial 1 | 9.0 (1.2)<sup>2</sup> | 10.0 (1.1) | 9.7 (1.1) | 6.2 (1.0) | 8.3 (0.5) | 7.2 (1.0) | 2.1 (1.2) | -1.1 (1.2) |
| Trial 2 | 8.6 (1.1) | 9.6 (0.4),<br>cast only-9.5 (0.3) | 9.4 (1.2)<br>cast only -<br>9.2 (0.9) | 6.5 (0.8) | 8.4 (0.1) | 8.1 (0.2) | 1.9 (0.9) | -0.3 (0.5) |
| Trial 3 | 8.5 (0.9) | 10.4 (0.8)<br>cast only-10.1 (0.9) | 9.7 (0.7) | 5.7 (0.6) | 9.1 (0.2) | 8.5 (0.3) | 3.4 (0.9) | -0.6 (0.7) |

<sup>1</sup>For the calculations cast, and non-cast sample estimates were combined since the values were not statistically different. Eight samples were analysed from each temperature treatment and each trial.
<sup>2</sup>Values in parentheses are standard deviations. The differences between warm and initial phase were significant ( $p < 0.008$ , Welch's *t*-test) for the three trials while the difference between recovery and warm samples was not significant for trial 1 or trial 2 and marginally significant for trial 3 ( $p = 0.03$ , Welch's *t*-test).
<sup>3</sup>Significantly different $p < 0.0006$ (Welch's *t*-test) between samples in each of the three trials.

### 3.5. Bacterial population estimation

Assuming 12 copies per cell and utilizing the proportions of Vibrionaceae in each sample (Supplementary Table S1) qPCR-based estimates for total 16S rRNA gene concentration lead to an approximation of cell number/g of fecal sample:

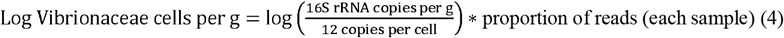

On this basis, the Vibrionaceae population was estimated to average 5.7 to 6.5 cells/g in the initial trial samples (Table 1). This population density increased to 8.3 to 9.1 log copies/g after the warm phase. In the recovery phase cell numbers were lower at 7.2 to 8.5 log copies/g. Overall, there was a significant (p<0.0006) increase in Vibrionaceae populations in the warm phase samples ranging on average from 1.9 to 3.4 log units (Table 1). Reducing the temperature in the recovery phase led to a significant reduction in Vibrionaceae abundance of 0.3 to 1.1 log units (Table 1).

## 4. Discussion

Numerous heat stress experiments involving fish have been recently performed to learn about the effects on microbiomes and overall health. Of these studies several involve commercially important salmonid species. The research designs vary with some exposing fish (typically hatchery smolt) to short term temperature effects (12 to 48 h, e.g. [42-44]). While others, such as a study on sturgeons [45] and chinook salmon [46] performed lengthier experiments (34 and 49 d, respectively) with larger smolt. In general heat stress alters the gut microbiomes. Heat stress studies performed in freshwater systems indicated increases in different bacterial genera, but certain commonalities can be observed. Sturgeon, rainbow trout, tsinling lenok trout, and chinook salmon all demonstrated increased gut microbiome presence of either or both *Cetobacterium* and *Aeromonas*-related species. For salmonids reared in seawater the natural seasonal effects of water temperature suggest Vibrionaceae being promoted under summer conditions [7, 13, 47-49]. Vibrionaceae also colonize wild salmon when they are at sea as observed in adults returning to rivers to spawn [18]. Predominance of Vibrionaceae can also be observed in other seawater farmed fish deliberately heat stressed, such as chum salmon [50]. Surveys of fish in estuaries also suggests higher temperatures drive increased presence of Vibrionaceae as well as increased dysbiosis [25].

In the experiments performed here Atlantic salmon were exposed to a temperature that was 3-6°C above their optimal temperature for growth [5] for an extended period of time. This is equivalent to an unusually long and warm summer period, mimicking heat waves previously experienced in the region [4]. In the Tasmanian region climate-based modelling has found that warmer summers would be detrimental to salmon farming productivity but could be offset by increased growth rates during corresponding warmer winter periods [51]. Since salmon take about 12 to 18 months to reach harvest weight the overall prospect of warmer sea surface temperatures is not ideal overall. Unlike other studies similar to this one we repeated the experiments three times and also performed qPCR to estimate bacterial population change overall. This was believed to be important since it is first unclear how reproducible microbiome studies are when dealing with different cohorts of fish. Secondly, the size of the microbial community in the gut of fish is often only vaguely defined. From these experiments we particularly wanted to determine the relationship of temperature exposure to categorized fecal consistency scores and cast production, another feature that other studies have not adopted for fish gut microbiome analysis though is prevalent in human studies [52].

In terms of fish associated impacts we observed that the tank warm phase resulted in a substantial increase in cast containing samples in the period when voluntary feeding had stopped (Fig. 2). The lack of voluntary feeding in salmon is recognized as a coping behavior owing to reduced health or environmental stress. Extended fasting leads to mortalities and size reduction and is considered a serious welfare issue [53]. We did not assess gut histology to determine if other impacts to gut health occur that might be contributing to voluntary feeding. It has been shown Atlantic salmon in sea cages experiencing high temperatures and reduced dissolve oxygen increase stress cortisol levels and develop increased intestinal hind gut epithelial layer permeability [54]. Furthermore, genes for cytolethal distending toxin and EAST1 heat-stable enterotoxin were detected in digesta samples from Atlantic salmon exposed to extended summer conditions in Tasmania [48]. The encoded proteins if produced at significant levels could lead to increased permeability in the gut along with suppressed local immune responses. It is unknown what bacterial effectors relate to cast production we can only suggest a temperature relationship so far. An increase in average fecal scores was observed in samples from Atlantic salmon collected during the summer (average 3.5 ± 0.1) in which sea surface temperatures reached 17-18°C [7]. By comparison, average scores in samples collected in winter (10-11°C) averaged only 2.0 ± 0.1 [7]. These corroborating results indicate casts can be another feature indicative of heat stressed Atlantic salmon. The use of repeated trials demonstrated that cast production varied. More cast only samples (score 4, Fig. 2) were obtained in trials 2 and 3 compared to trial 1. The reasons for this remain unclear but we hypothesize this could reflect the predominant bacterial species proliferating in samples during the different trials (discussed below). To understand this further we examined the species composition as well as changes in populations levels.

By using qPCR and applying total 16S rRNA gene abundance as a reference frame [55] it was possible to estimate approximate Vibrionaceae populations in digesta and cast samples after each of the temperature phases. To estimate bacterial populations there is a need to account for the contribution of chloroplast and mitochondria derived 16S rRNA genes of which the vast majority are from feed plant components. Plant chloroplast and mitochondria genome number vary with plant maturity [56] plus it is assumed the feed contains also diverse reads from bacteria as noted by Karlsen and colleagues [14]. Thus, calculating 16S rRNA copy numbers to estimate the feed carry over is challenging without using specific primer sets. However, feed DNA likely being from inactive organisms (since feed pellets are subjected to wet heat of 100°C for 20 minutes during manufacture) can be assumed to only decrease in the fecal samples due to dilution and digestion [13]. Thus, for making estimations we simply ignored the feed background and calculated the approximate Vibrionaceae population given it makes up the largest proportion of reads overall. *Vibrio* and *Aliivibrio* bacterial species have 10 to 13 16S rRNA gene copies per genome (rrnDB v. 5.9 [57]). The analysis includes the assumption that the PCR process used for generating sequences is not biased, affecting the proportional representation in a systematic way. We do not think this is an issue since the replicated trials provide a degree of understanding of the experimental variation. Also, the goal was more to determine the overall abundance differences as opposed to precise absolute abundance estimates for specific samples.

From the results it was clearly observed that most stripped fecal samples from the warm phase became highly enriched in members of family Vibrionaceae (Fig. 5 and 6). This was clearly due to growth occurring in digesta and cast-containing samples (Table 1). The population expansion resulted in much lower alpha diversity (Fig. 3). The overall responses had the same pattern between all three trials based on qPCR data and cell count estimates. The Vibrionaceae proliferation occurred during the warm phase. When the tank water temperatures were reduced 3.5-4°C to 15°C there was a small but significant decline in the Vibrionaceae population (Table 1). This response can simply put down to growth biokinetics [58]. The warm phase temperature resulted in more rapid growth rates while reducing the temperature reduces the growth rate of Vibrionaceae subsequently affecting how much biomass can accumulate in tandem with fish feeding and excretion.

We have hypothesized previously Vibrionaceae, in particular *Aliivibrio* are a cause of the production of fecal casts in fish sampled during summer [7]. Fecal cast production in finfish has been attributed to disease, stress, or simply the lack of feeding [59-61]. The expected higher growth rates and biomass of *Aliivibrio* and *Vibrio* spp. during warm summers may increase the risk of dysbiosis or disease in farmed salmon, but this relies on strains directly or indirectly affecting the host negatively. The experiments indicated only specific Vibrionaceae respond strongly to warm water conditions in a given trial alongside a high incidence in fecal cast production. Since distinct species predominated in each of the three trials performed this suggests that fish may be colonized successfully by a number of species. Further cast production may not be a species-specific effect but could be generalized response to bacterial overgrowth. Even then at the current stage of knowledge a rigorously provable relationship can be established that specific Vibrionaceae are responsible for influencing fecal cast production or that affect feeding rates.

The primary proliferating OTUs from trial 1 and 2 heat stress were classified as *Vibrio scophthalmi*. This species is considered an opportunistic pathogen of farmed fish [62-65] but in most situations it is more likely to be a harmless commensal [66]. Overgrowth of these OTUs only affected the fish concerned temporarily in terms of feeding, since they resumed feeding in the recovery phase. This species has been detected at considerable abundance in seawater farmed Atlantic salmon [7, 13, 17] and thus seems a well-adapted colonizing species. Genome-based analysis is needed to ascertain whether the associated *V. scophthalmi* OTUs constitute one or more species and also enable determination of whether they have any potential for virulence.

In trial 2 *Aliivibrio* OTUs also grew, eventually becoming predominant after the recovery phase. Based on CLR averages this increase amounted to approximately 3.7 log units on average. The OTUs of this population are related to the *Aliivibrio sifiae* group with sequences the same as *Aliivibrio* observed in the study of Bozzi et al. [20]. This group includes several closely related species [57] and includes the causative agent of cold water vibriosis - *Aliivibrio salmonicida*. From trial 2 the occurrence of cast containing samples remained at the same frequency in the recovery phase as the warm phase. This is unlike the reduction observed for trials 1 and 3 (Fig. 2). Whether the *Aliivibrio* strains present in Trial 2 exert a potentially enhanced dysbiotic influence, resulting in this higher proportion of cast only samples, remains to be elucidated. In the study of Bozzi et al. [20] related *Aliivibrio* strains were considered to be opportunistically pathogenic. These OTUs became more abundant in gut samples in unhealthy fish suffering from a skin infection caused by *Tenacibaculum dicentrarchi*. This group of *Aliivibrio* seem common to Atlantic salmon and based on distribution seem to grow well under cooler conditions [67]. Further research is needed to determine if the incidences of this *Aliivibrio* group is frequent enough to be of concern and can be observed impacting fish health when in combination with other stressors or infections. Since we only observed the *A. sifiae* group strains in one of three trials and then only at elevated levels at the lower tested temperature this may suggest this group could be less of a concern under heat stress.

In trial 3 the main species proliferating was *Aliivibrio finisterrensis* [68]. The biology of this species is currently poorly known though it has been observed to be one of the dominant Vibrionaceae in Atlantic salmon and Chinook salmon farmed in seawater in both Tasmania and New Zealand [7, 14, 17, 49]. Typically, in seawater salmon it is predominant in fish alongside *Vibrio scophthalmi*. This suggests this species group are also good colonizers of the Atlantic salmon gut. The results suggest the growth seems to be stimulated under warmer conditions, potentially leading to overgrowth.

The negative effects of the overgrowth of colonizing bacteria have been discussed in relation to dysbiosis and disease [69]. However, specific experiments are needed to understand what the thresholds for populations in the gut of Atlantic salmon leading to observable negative health impacts and also how this relates to aspects of fish biology. Besides the occurrence of casts and reduced feeding none of aforementioned OTUs seem to be invasive, acknowledging the high populations they can reach. *Aliivibrio salmonicida* for example was a major disease problem for Norwegian salmon aquaculture until vaccines were developed [70, 71]. This species pathogenesis involves rapid invasiveness and tissue damage due to an endotoxic lipopolysaccharide [72], chitinolytic enzymes that enable spread in the gut [23], and other unknown effectors. Besides this species and the faunal mutualist *Aliivibrio fischeri* [73] other *Aliivibrio* species have not been investigated in any detail. The high populations of Vibrionaceae in cast-containing samples (as much as log 9 cells/g) suggests various species can form dense biofilms on and in the Atlantic salmon gut mucosal layer [74]. This has been studied most extensively in relation to *Vibrio cholerae* forming biofilms in the human gastrointestinal tract [75]. Predominant species occurring within the three trials could be due to exertion of competitive exclusion [76] against other possible bacterial colonists. The exclusion comes about owing to the successful species being already quite abundant in the initial samples. The initial Vibrionaceae populations were estimated to be in the range of 6 log units/g feces (Table 1). The OTUs we detected could be evolved to form biofilm naturally in the gut of Atlantic salmon and other fish species due to possessing inherent antagonism capabilities and fitness advantages. Bacterial strains showing evolved biofilm formation mechanisms may also potentially trade this capability for reduced virulence [77]. This latter general concept could explain why the high Vibrionaceae populations occurring in the Atlantic salmon gut is not equated with obvious disease states.

Microbiome changes induced by environmental stressors and determinants (temperature, salinity, pH, O2, photoperiod length) have been suggested to potentially impact gut mucosal immunity [78]. Stressors could induce increased responses as described by Sundh et al. [54] when overgrowth of gut bacteria occurs at the same time. Further research connecting immune system functions to heat stress and microbiome function is necessary to clarify the relationships and significance to Atlantic salmon health, especially when under sustained warm conditions. This would also be aided by developing an understanding of the mechanisms leading to cast excretion.

## Conclusions

The results of the repeated heat stress trials demonstrate that cast excretion is accompanied by increased predominance of Vibrionaceae. This increase results from population growth in the range of 2 to 3 log units and could be observed quantitatively occurring in both digesta and in cast samples. The predominant species that proliferate under warm conditions differed between the three trials performed but included a narrow diversity of bacteria including *Vibrio scophthalmi*, and the *Aliivibrio sifiae* and *Aliivibrio finisterrensis* species groups. The role these species play in inducing cast excretion requires more elucidation but due to their high populations in cast material are ideal candidates for further study. Such efforts would be well supported by also obtaining further developing understanding of the impact of environmental stressors, in particular increased temperature, on gut mucosal physiology and immunology in Atlantic salmon. These studies would need to include functional interrelations with the gut microbiome.

## Supplementary Materials

The following supporting information can be downloaded at: www.mdpi.com/xxx/s1, Table S1. OTU read table for the three temperature treatment trials.

## Institutional Review Board Statement

All animal care and experimental procedures in this study were authorized under University of Tasmania Animal Ethics permits A0015452 (approved 5th April 2015), A0016275 (approved 7th February 2016), and A0017573 (approved 7th May 2017).

## Informed Consent Statement

not applicable.

## Data Availability Statement

The raw 16S rRNA gene sequence data and metadata files are deposited in the NCBI SRA database under BioProject codes: trial 1 (PRJNA1023021), trial 2 (PRJNA1023026), and trial 3 (PRJNA1023294). Other data is available on request to the corresponding author.

## Acknowledgments

The authors would like to thank staff at Taroona EAF (Institute of Marine and Antarctic Studies, University of Tasmania) for tank set up, regular feeding, and monitoring.

## References

1. Hevrøy, E.M.; Hunskår, C.; De Gelder, S.; Shimizu, M.; Waagbø, R.; Breck, O.; Takle, H.; Sussort, S.; Hansen, T., GH–IGF system regulation of attenuated muscle growth and lipolysis in Atlantic salmon reared at elevated sea temperatures. J. Comp. Physiol. B. 2012, 183, 243–259. 10.1007/s00360-012-0704-5.

2. Strom, J.F.; Thorstad, E.B.; Rikardsen, A.H. Thermal habitat of adult Atlantic salmon Salmo salar in a warming ocean. J. Fish Biol. 2020, 96, 327–336. 10.1111/jfb.14187.

3. Debes, P.V.; Solberg, M.F.; Matre, I.H.; Dyrhovden, L.; Glover KA.; Genetic variation for upper thermal tolerance diminishes within and between populations with increasing acclimation temperature in Atlantic salmon. Heredity (Edinb) (2021) 127(5), 455–466. 10.1038/s41437-021-00469-y.

4. Wade, N.M.; Clark, T.D.; Maynard, B.T.; Atherton, S.; Wilkinson, R.J.; Smullen, R.P.; Taylor, R.S. Effects of an unprecedented summer heatwave on the growth performance, flesh colour and plasma biochemistry of marine cage-farmed Atlantic salmon (Salmo salar). J. Thermal Biol. 2019, 80, 64–74. 10.1016/j.jtherbio.2018.12.021.

5. Calado, R.; Mota, V.C.; Madeira, D.; Leal, M.C. Summer is coming! Tackling ocean warming in Atlantic Salmon cage farming. Animals (Basel). 2021, 11, 1800. 10.3390/ani11061800.

6. Grünenwald, M.; Adams, M. B.; Carter, C. G.; Nichols, D. S.; Koppe, W.; Verlhac-Trichet, V.; Schierle, J.; Adams, L.R. Pigment-depletion in Atlantic salmon (Salmo salar) post-smolt starved at elevated temperature is not influenced by dietary carotenoid type and increasing α-tocopherol level. Food Chem. 2019, 299, 125140. 10.1016/j.foodchem.2019.125140.

7. Reid, C.E.; Bissett, A.; Huynh, C.; Bowman, J.P.; Taylor, R.S. Time from feeding impacts farmed Atlantic salmon (Salmo salar) gut microbiota and fecal score. Aquaculture 2024, 579, 740174. 10.1016/j.aquaculture.2023.740174.

8. Hudson, J.; Adams, M.; Jantawongsri, K.; Dempster, T.; Nowak, B.F; Evaluation of low temperature and salinity as a treatment of Atlantic salmon against amoebic gill disease. Microorganisms. 2022, 10, 202. 10.3390/microorganisms10020202.

9. Soto, D.; León-Muñoz, J.; Garreaud, R.; Quiñones, R.A.; Morey, F.; Scientific warnings could help to reduce farmed salmon mortality due to harmful algal blooms. Marine Policy. 2021, 132, 104705. 10.1016/j.marpol.2021.104705.

10. Bravo, S.; Treasurer, J. The management of the sea lice in Chile: A review. Rev. Aquacultur. 2023, 15, 1749–1764. 10.1111/raq.12815.

11. Burke, M.; Grant, J.; Filgueira, R.; Stone T. Oceanographic processes control dissolved oxygen variability at a commercial Atlantic salmon farm: Application of a real-time sensor network. Aquaculture. 2021, 533, 736143. 10.1016/j.aquaculture.2020.736143.

12. Kim, P.S.; Shin, N.R.; Lee, J.B.; Kim, M.S.; Whon, T.W.; Hyun, D.W.; Yun, J.H.; Jung, M.J.; Kim, J.Y.; Bae, J.W. Host habitat is the major determinant of the gut microbiome of fish. Microbiome. 2021, 9, 166. 10.1186/s40168-021-01113-x.

13. Zarkasi, K.Z.; Abell, G.C.; Taylor, R.S.; Neuman, C.; Hatje, E.; Tamplin, M.L.; Katouli, M.; Bowman, J.P. Pyrosequencing-based characterization of gastrointestinal bacteria of Atlantic salmon (Salmo salar L.) within a commercial mariculture system. J. Appl. Microbiol. 2014, 117, 18–27. 10.1111/jam.12514.

14. Karlsen, C.; Tzimorotas, D.; Robertsen, E.M.; Kirste, K.H.; Bogevik, A.S.; Rud, I. Feed microbiome: confounding factor affecting fish gut microbiome studies. ISME Commun. 2022, 2, 14. 10.1038/s43705-022-00096-6.

15. Lorgen-Ritchie, M.; Clarkson, M.; Chalmers, L.; Taylor, J.F.; Migaud, H.; Martin, S.A.M. A temporally dynamic gut microbiome in Atlantic Salmon during freshwater recirculating aquaculture system (RAS) production and post-seawater transfer. Front. Marine Sci. 2021, 8, 711797. 10.3389/fmars.2021.711797.

16. Wang, J.; Jaramillo-Torres, A.; Li, Y.; Kortner, T.M.; Gajardo, K.; Brevik, Ø.J.; Jakobsen, J.V.; Krogdahl, Å. Microbiota in intestinal digesta of Atlantic salmon (Salmo salar), observed from late freshwater stage until one year in seawater, and effects of functional ingredients: a case study from a commercial sized research site in the Arctic region. Animal Microbiome. 2021, 3, 14. 10.1186/s42523-021-00075-7.

17. Zarkasi, K.Z.; Taylor, R.S.; Glencross, B.D.; Abell, G.C.J.; Tamplin, M.L.; Bowman, J.P. Atlantic Salmon (Salmo salar L.) gastrointestinal microbial community dynamics in relation to digesta properties and diet. Microb. Ecol. 2016, 71, 589–603. 10.1007/s00248-015-0728-y.

18. Llewellyn, M.; McGinnity, P.; Dionne, M.; Letourneau, J.; Thonier, F.; Carvalho, G.R.; Creer, S.; Derome, N.; The biogeography of the Atlantic salmon (Salmo salar) gut microbiome. ISME J. 2016, 10, 1280–1284. 10.1038/ismej.2015.18.

19. Rasmussen, J.A.; Killerich, P.; Madhun, A.S.; Waagbø, R.; Lock, E.J.R.; Madsen, L.; Gilbert, M.T.P.; Kristiansen, K.; Limborg, M.T. Co-diversification of an intestinal Mycoplasma and its salmonid host. ISME J. 2023, 17, 682– 692. 10.1038/s41396-023-01379-z.

20. Bozzi, D.; Rasmussen, J.A.; Carøe, C.; Sveier, K.; Nordøy, K.; Gilbert, M.T.P.; Limborg, M.T. Salmon gut microbiota correlates with disease infection status: potential for monitoring health in farmed animals. Animal Microbiome. 2021, 3, 30. 10.1186/s42523-021-00096-2.

21. López, J.R.; Lorenzo, L.; Alcantara, R.; Navas, J.I. Characterization of Aliivibrio fischeri strains associated with disease outbreak in brill Scophthalmus rhombus. Dis. Aquat. Organ. 2017, 124, 215–222. 10.3354/dao03123.

22. Baker-Austin, C.; Oliver, J.D.; Alam, M.; Ali, A.; Waldor, M.K.; Qadri, F.; Martinez-Urtaza, J. Vibrio spp. infections. Nature Rev. Dis. Primers. 2018, 4, 8. 10.1038/s41572-018-0005-8.

23. Skåne, A.; Edvardsen, P.K.; Cordara, G.; Loose, J.S.M.; Leitl, K.D.; Krengel, U.; Sørum, H.; Askkarian, F.; Vaaje-Kolstad, G. Chitinolytic enzymes contribute to the pathogenicity of Aliivibrio salmonicida LFI1238 in the invasive phase of cold-water vibriosis. BMC Microbiol. 2022, 22, 194. 10.1186/s12866-022-02590-2.

24. Klakegg, Ø.; Myhren, S.; Juell, R.A.; Aase, M.; Salonius K.; Sørum H. Improved health and better survival of farmed lumpfish (Cyclopterus lumpus) after a probiotic bath with two probiotic strains of Aliivibrio. Aquaculture. 2020, 518, 734810. 10.1016/j.aquaculture.2019.734810.

25. Suzzi, A.L.; Stat, M.; Gaston, T.F.; Siboni, N.; Williams, N.L.R.; Seymour, J.; Huggett, M.J. Elevated estuary water temperature drives fish gut dysbiosis and increased loads of pathogenic Vibrionaceae. Environ. Res. 2023, 219, 115144.

26. Oliver, E.C.; Benthuysen, J.A.; Bindoff, N.L.; Hobday, A.J.; Holbrook, N.J.; Mundy, C.N.; Perkin-Kirkpatrick, S.E. The unprecedented 2015/16 Tasman Sea marine heatwave. Nature Commun. 2017, 8, 16101. 10.1038/ncomms16101.

27. Foddai, M.; Carter, C.G.; Hilder, P.E.; Gurr, H.; Ruff, N. Combined effects of elevated rearing temperature and dietary energy level on heart morphology and growth performance of Tasmanian Atlantic salmon (Salmo salar L.). J. Fish Dis. 2022, 45, 301–313. 10.1111/jfd.13555.

28. Lane, D.J. 16S/23S rRNA sequencing. In Nucleic acid techniques in bacterial systematics. Stackebrandt, E.; Goodfellow M., Eds.; Wiley: New York City, New York, USA, 1991; pp. 115–175. 10.1002/jobm.3620310616.

29. Turner, S.; Pryer, K.M.; Miao, V.P.W.; Palmer, J.D. Investigating deep phylogenetic relationships among cyanobacteria and plastids by small subunit rRNA sequence analysis. J. Euk. Biol. 1999, 46, 327–338. 10.1111/j.1550-7408.1999.tb04612.x.

30. Callahan, B.J.; McMurdie, P.J.; Rosen, M.J.; Han, A.W.; Johnson, A.J.; Holmes, S.P. DADA2: High-resolution sample inference from Illumina amplicon data. Nat. Methods. 2016, 13, 7, 581–583. 10.1038/nmeth.3869.

31. Rognes, T.; Flouri, T.; Nichols, B.; Quince, C.; Mahé, F. VSEARCH: a versatile open source tool for metagenomics. Peer J. 2016, 4, e2584. 10.7717/peerj.2584.

32. Bolyen, E.; Riseout, J.R.; Dillon, M.R.; Bokulich, N.A.; Abnet, C.C.; Al-Ghalith, G.A.; Alexander, H.; Alm, E.J.; Arumugam, M.; Asnicar, F.; et al. Reproducible, interactive, scalable and extensible microbiome data science using QIIME 2. Nat. Biotechnol. 2019, 37, 852–857. 10.1038/s41587-019-0252-6.

33. Quast, C.; Pruesse, E.; Yilmaz, P.; Gerken, J.; Schweer, T.; Yarzo, P.; Peplies, J.; Glöckner, F.O.; The SILVA ribosomal RNA gene database project: improved data processing and web-based tools. Nucl. Acids Res. 2013, 41, D590–596. 10.1093/nar/gks1219.

34. Davis, N.M.; Proctor, D.M.; Holmes, S.P.; Relman, D.A.; Callahan, B.J.; Simple statistical identification and removal of contaminant sequences in marker-gene and metagenomics data. Microbiome 2018, 6, 226. 10.1186/s40168-018-0605-2.

35. Aitchison, J. The Statistical Analysis of Compositional Data. Chapman and Hall, London; New York; 1982. 10.1111/j.2517-6161.1982.tb01195.x.

36. Quinn, T.P.; Erb. I.; Gloor, G.; Notredame, C.; Richardson, M.F.; Crowley, T.M. A field guide for the compositional analysis of any-omics data. Gigascience. 2019, 8, giz107. 10.1093/gigascience/giz107.

37. Anderson, M.J.; Willis, T.J. Canonical analysis of principal coordinates: a useful method of constrained ordination for ecology. Ecology. 2003, 84, 511–525. 10.1890/0012-9658(2003)084[0511:CAOPCA]2.0.CO;2.

38. Ratkowsky, D.A. Choosing the number of principal coordinates when using CAP, the canonical analysis of principal coordinates. Austral Ecol. 2016, 41, 842–851 10.1111/aec.12378.

39. Anderson, M.J. Permutational multivariate analysis of variance (PERMANOVA). Wiley StatsRef: Statistics Reference Online; 2017. 10.1002/9781118445112.stat07841.

40. Suzuki, M.T.; Taylor, L.T.; DeLong, E.F. Quantitative analysis of small-subunit rRNA genes in mixed microbial populations via 5’-nuclease assays. Appl. Environ. Microbiol. 2000, 66, 4605–4614. 10.1128/AEM.66.11.4605-4614.2000.

41. Bustin, S.A.; Benes, V.; Garson, J.A.; Hellemans, J.; Huggett, J.; Kubista, M.; Mueller, R.; Nolan, T.; Pfaffl, M.W.; Shipley, G.L.; et al. The MIQE guidelines: minimum information for publication of quantitative real-time PCR experiments. Clin. Chem. 2009, 55, 4, 611-622. 10.1373/clinchem.2008.112797.

42. Fang, M.; Lei, Z.; Ruilin, M.; Jing, W.; Leqiang, D. Hot temperature stress induced oxidative stress, gut inflammation and disordered metabolome and microbiome in tsinling lenok trout. Ecotox. Environ. Safety. 2023, 266, 115607. 10.1016/j.ecoenv.2023.115607

43. Zhou, C.; Yang, S.; Ka, W.; Gao, P.; Li, Y., Long, R.; Wang J. Association of gut microbiota with metabolism in rainbow trout under acute heat stress. Front. Microbiol. 2022, 13, 846336. 10.3389/fmicb.2022.846336.

44. Zhao, Z.; Zhao, H.; Wang, X.; Zhang, L.; Mou, C.; Huang, Z.; Ke, H.; Duan, Y.; Zhou, J.; Li, Q. Effects of different temperatures on Leiocassis longirostris gill structure and intestinal microbial composition. Sci. Rep. 2024, 14, 7150. 10.1038/s41598-024-57731-6.

45. Yang, S.; Zhang, C; Xu, W; Li, D; Feng, Y; Wu, J; Luo, W; Du, X; Du, Z.; Huang, X. Heat stress decreases intestinal physiological function and facilitates the proliferation of harmful intestinal microbiota in sturgeons. Front. Microbiol. 2022, 13, 755369. 10.3389/fmicb.2022.755369.

46. Steiner, K.; Laroche, O.; Walker, S.P.; Symonds, J.E. Effects of water temperature on the gut microbiome and physiology of Chinook salmon (Oncorhynchus tshawytscha) reared in a freshwater recirculating system. Aquaculture 2022, 560, 738529. 10.1016/j.aquaculture.2022.738529

47. Huyben, D.; Sun, L.; Moccia, R.; Kiessling, A.; Dicksved, J; Lundh, T. Dietary live yeast and increased water temperature influence the gut microbiota of rainbow trout. J. Appl. Microbiol. 2018, 124, 1377–1392. 10.1111/jam.13738.

48. Neuman, C.; Hatje, E.; Zarkasi, K.Z.; Smullen, R.; Bowman, J.P.; Katouli, M. The effect of diet and environmental temperature on the faecal microbiota of farmed Tasmanian Atlantic Salmon (Salmo salar L.). Aquac Res. 47, 660–672. 10.1111/are.12522.

49. Zhao, R.; Symonds, J.E.; Walker, S.; Steiner, K.; Carter, C.G.; Bowman, J.P.; Nowak, B.F.; Salinity and fish age affect the gut microbiota of farmed Chinook salmon (Oncorhynchus tshawytscha). Aquaculture. 2020, 528, 735539. 10.1016/j.aquaculture.2020.735539.

50. Ghosh SK, Wong MK-S, Hyodo S, Goto S and Hamasaki K (2022) Temperature modulation alters the gut and skin microbial profiles of chum salmon (Oncorhynchus keta). Front. Mar. Sci. 9:1027621. 10.3389/fmars.2022.1027621.

51. Meng, H.; Hayashida, H.; Norazmi-Lokman, N.H.; Strutton, P.G. Benefits and detrimental effects of ocean warming for Tasmanian salmon aquaculture. Cont. Shelf Res. 2022, 246, 104829. 10.1016/j.csr.2022.104829. 10.1016/j.csr.2022.104829.

52. Vandeputte, D.; Falony, G.; Vieira-Silva, S.; Tito, R.Y.; Joossens, M.; Raes, J. Stool consistency is strongly associated with gut microbiota richness and composition, enterotypes and bacterial growth rates. Gut. 2016, 65, 57–62. 10.1136/gutjnl-2015-309618.

53. Hvas, M.; Kolarevic, J.; Noble, C.; Oppedal, F.; Stien, L.H. Fasting and its implications for fish welfare in Atlantic salmon aquaculture. Rev. Aquac., 2024, 16, 1–25. 10.1111/raq.12898.

54. Sundh, H.; Kvamme, B.O.; Fridell, F.; Olsen, R.E.; Ellis, T.; Taranger, G.L.; Sundell, K. Intestinal barrier function of Atlantic salmon (Salmo salar L.) post smolts is reduced by common sea cage environments and suggested as a possible physiological welfare indicator. BMC Physiol. 2010, 10, 22. 10.1186/1472-6793-10-22.

55. Morton, J.T.; Marotz, C.; Washburne, A.; Silverman, J.; Zaramela, L.S.; Edlund, A.; Zengler, K.; Knight, R. Establishing microbial composition measurement standards with reference frames. Nat. Commun. 2019, 10, 2719. 10.1038/s41467-019-10656-5.

56. Kubínová, Z.; Janácek, J.; Lhotáková, Z.; Kubínová, L.; Albrechtová, J. Unbiased estimation of chloroplast number in mesophyll cells: advantage of a genuine three-dimensional approach. J. Exp. Bot. 2014, 65, 609–620. 10.1093/jxb/ert407.

57. Stoddard, S.F, Smith, B.J.; Hein, R.; Roller, B.R.K.; Schmidt, T.M. rrnDB: improved tools for interpreting rRNA gene abundance in bacteria and archaea and a new foundation for future development. Nucl. Acids Res. 2015, 43, D595–D598. 10.1093/nar/gku1201.

58. Corkrey, R.; McMeekin, T.A.; Bowman, J.P.; Ratkowsky, D.A., Olley, J.; Ross, T. The biokinetic spectrum for temperature. PLOS ONE. 2016, 11, e0153343. 10.1371/journal.pone.0153343.

59. Turner Jr., J.W.; Nemeth, R.; Rogers, C. Measurement of fecal glucocorticoids in parrotfishes to assess stress. Gen. Comp. Endocrinol. 2003, 133, 341–352. 10.1016/S0016-6480(03)00196-5.

60. Dhar, A.K.; LaPatra, S.; Orry, A.; Allnutt, F.C.T.; Infectious pancreatic necrosis virus. In: Fish viruses and bacteria: pathobiology and protection. Woo, P.T.K.; Cipriano, T.C., Eds. CABI Digital Library; 2017, pp. 1–13. 10.1079/9781780647784.00

61. Waagbø, R.; Jørgensen, S.M.; Timmerhaus, G.; Breck, O.; Olsvik, P.A. Short-term starvation at low temperature prior to harvest does not impact the health and acute stress response of adult Atlantic salmon. Peer J. 2017, 5, e3273. 10.7717/peerj.3273.

62. Zhang, Z.; Yu, Y-X.; Wang, Y-G.; Liu, X.; Wang, L-F.; Zhang, H.; Liao, M-J.; Li, B. Complete genome analysis of a virulent Vibrio scophthalmi strain VSc190401 isolated from diseased marine fish half-smooth tongue sole, Cynoglossus semilaevis. BMC Microbiol. 2020, 20, 341. 10.1186/s12866-020-02028-7.

63. Yu, Y.; Liu, X.; Wang, Y.; Liao, M.; Tang, M.; Rong, X.; Wang, C.; Li, B.; Zhang, Z. Antimicrobial resistance and genotype characteristics of Vibrio scophthalmi isolated from diseased mariculture fish intestines with typical inter-annual variability. Front. Mar. Sci. 2022, 9, 924130. 10.3389/fmars.2022.924130

64. Ayala, A.J.; Ogbunugafor, C.B. When vibrios take flight: A meta-analysis of pathogenic Vibrio species in wild and domestic birds. Adv. Exp. Med. Biol. 2023, 1404, 295–336. 10.1007/978-3-031-22997-8_15. PMID: 36792882.

65. Liu, X.; You, C.; Zeng, Y.; Isolation and identification of pathogenic Vibrio species in black rockfish Sebastes schlegeli. Fishes. 2023, 8, 235. 10.3390/fishes8050235.

66. García-Aljaro, C.; Melado-Rovira, S.; Milton, D.L.; Blanch, A.R.; Quorum-sensing regulates biofilm formation in Vibrio scophthalmi. BMC Microbiol. 2012, 12, 287. 10.1186/1471-2180-12-287.

67. Klemetsen, T.; Karlsen, C.R.; Willassen, NP. Phylogenetic revision of the genus Aliivibrio: intra- and inter-species variance among clusters suggest a wider diversity of species. Front. Microbiol. 2021, 12, 626759. 10.3389/fmicb.2021.626759.

68. Beaz-Hidalgo, R.; Doce, A.; Balboa, S.; Barja, J.L.; Romalde, J.L. Aliivibrio finisterrensis sp. nov.; isolated from Manila clam, Ruditapes philippinarum and emended description of the genus Aliivibrio. Int. J. Syst. Evol. Microbiol. 2010, 60, 223–228. 10.1099/ijs.0.010710-0.

69. Zhao, R.; Symonds, J.E.; Walker, S.P.; Steiner, K.; Carter, C.G.; Bowman, J.P.; Nowak, B.F. Relationship between gut microbiota and Chinook salmon (Oncorhynchus tshawytscha) health and growth performance in freshwater recirculating aquaculture systems. Front. Microbiol. 2023, 14, 1065823. 10.3389/fmicb.2023.1065823.

70. Hjerde, E.; Lorentzen, M.S.; Holden, M.T.; Seeger, K.; Paulsen, S.; Bason, N.; Churcher, C.; Harris, D.; Norbertczak, H.; Quail, M.A.; et al. The genome sequence of the fish pathogen Aliivibrio salmonicida strain LFI1238 shows extensive evidence of gene decay. BMC Genomics. 2008, 9, 616. 10.1186/1471-2164-9-616.

71. Kashulin, A.; Seredkina, N.; Sørum H. Cold-water vibriosis. The current status of knowledge. J. Fish Dis. 2017, 40, 119–126. 10.1111/jfd.12465.

72. Nørstebø, S.F.; Lotherington, L.; Landsverk, M.; Bjelland, A.M.; Sørum, H. Aliivibrio salmonicida requires O-antigen for virulence in Atlantic salmon (Salmo salar L.). Microb. Pathogens. 2018, 124, 322–331. 10.1016/j.micpath.2018.08.058.

73. Fung, B.L.; Esin, J.J.; Visick, K.L.; Vibrio fischeri: a model for host-associated biofilm formation. J. Bacteriol. 2024, 206, e0037023. 10.1128/jb.00370-23.

74. Jandl, B.; Dighe, S.; Baumgartner, M.; Makristathis, A.; Gasche, C.; Muttenthaler, M.; Gastrointestinal biofilms: endoscopic detection, disease relevance, and therapeutic strategies. Gastroenterol. 2024 article in press 12 June 2024. 10.1053/j.gastro.2024.04.032

75. Cho, J.Y.; Liu, R.; Macbeth, J.C.; Hsiao, A. The interface of Vibrio cholerae and the gut microbiome. Gut Microbes. 2021, 13, 1937015. 10.1080/19490976.2021.1937015.

76. Rendueles, O.; Ghigo, J-M. Mechanisms of competition in biofilm communities. Microbiol. Spectrum. 2015, 3, MB-0009-2014. 10.1128/microbiolspec.MB-0009-2014.

77. Tang, M.; Yang, R.; Zhuang, Z.; Han, S.; Sun, Y.; Li, P.; Fan, K.; Cai, Z.; Yang, Q.; Yu, Z.; et al. Divergent molecular strategies drive evolutionary adaptation to competitive fitness in biofilm formation. ISME J. 2024, 18, wrae135. 10.1093/ismejo/wrae135.

78 Morshed, S.M.; Lee. T.H. The role of the microbiome on fish mucosal immunity under changing environments. Fish Shellfish Immunol. 2023, 139,108877. 10.1016/j.fsi.2023.108877

